# Phosphorylation of mitochondrial matrix proteins regulates their selective mitophagic degradation

**DOI:** 10.1101/513135

**Authors:** Panagiota Kolitsida, Jianwen Zhou, Michal Rackiewicz, Vladimir Nolic, Jörn Dengjel, Hagai Abeliovich

**Affiliations:** Dept. of Biochemistry, Food Science and Nutrition, Hebrew University of Jerusalem, Rehovot, Israel; Department of Biology, University of Fribourg, Chemin du Musée 10, 1700 Fribourg, Switzerland; Department of Dermatology, Medical Center - University of Freiburg, Hauptstr. 7, 79104 Freiburg, Germany; Freiburg Institute for Advanced Studies, University of Freiburg, Albertstr. 19, 79104 Freiburg, Germany

**Author notes:** Address correspondence to: Dr. Hagai Abeliovich, Dept. of Biochemistry and Food Science, Hebrew University of Jerusalem, Rehovot, Israel 76100.

**Keywords:** mitophagy, autophagy, mitochondria, phosphorylation, phosphatase, Saccharomyces cerevisiae

## Abstract

Mitophagy is an important quality control mechanism in eukaryotic cells, and defects in mitophagy correlate with aging phenomena and neurodegenerative disorders. It is known that different mitochondrial matrix proteins undergo mitophagy with very different rates, but to date the mechanism underlying this selectivity at the individual protein level has remained obscure. We now present evidence indicating that protein phosphorylation within the mitochondrial matrix plays a mechanistic role in regulating selective mitophagic degradation in yeast, via involvement of the Aup1 mitochondrial protein phosphatase, as well as two known matrix-localized protein kinases, Pkp1 and Pkp2. By focusing on a specific matrix phosphoprotein reporter, we also demonstrate that phospho-mimetic and non-phosphorylatable point mutations at known phosphosites in the reporter increased or decreased its tendency to undergo mitophagy. Finally, we show that phosphorylation of the reporter protein is dynamically regulated during mitophagy, in an Aup1-dependent manner. Our results indicate that structural determinants on a mitochondrial matrix protein can govern its mitophagic fate, and that protein phosphorylation regulates these determinants.

**Significance statement:** Mitochondrial dysfunction underlies many age-related human pathologies. In normal cells, defective mitochondria are often degraded by mitophagy, a process in which these mitochondria are engulfed in autophagosomes and sent for degradation in the lysosome/vacuole. Surprisingly, studies on mitophagy in diverse eukaryotic organisms reveal an unexpected dimension of protein-level selectivity, wherein individual protein species exhibit divergent rates of mitophagic degradation. In this manuscript, we show that this surprising intra-mitochondrial selectivity can be generated by differential phosphorylation of individual mitochondrial protein species, and we identify mitochondrial phosphatases and kinases which contribute to this regulation. By identifying a mechanism which regulates the intra-mitochondrial selectivity of mitophagic degradation, our findings open the door to potential manipulation of the quality control process in the future.

## Introduction

Mitochondrial autophagy, or mitophagy, is considered an important quality control mechanism in eukaryotic cells (1–3). Defects in mitophagic clearance of malfunctioning mitochondria have been proposed to play a role in the pathogenesis of neurological disorders such as Parkinson’s, Alzheimer’s and Huntington’s diseases (4–8) and may be associated with additional aging-related pathologies (9). As with other forms of selective autophagy, mitophagy is induced by the activation of a receptor protein, or by its recruitment to the organellar surface. The receptor interacts with the autophagic machinery to mediate the engulfment of specific mitochondria which are designated for degradation (10, 11). In many mammalian cell types, loss of mitochondrial membrane potential stabilizes the PINK1 protein kinase on the outer mitochondrial membrane (12). This leads to the PINK1-dependent recruitment and phosphorylation of Parkin, a ubiquitin E3 ligase (13–15), as well as to the local production of phospho-ubiquitin (16–19). Phospho-ubiquitylation of mitochondrial outer membrane proteins recruits the soluble mitophagy receptors NDP52 and optineurin, which link the defective mitochondrial compartment with the autophagic machinery and mediate its degradation (20). Mitophagy has also been implicated in specific developmental transitions in mammals, such as muscle, neuron, and erythrocyte differentiation (21–25). In yeast cells, the Atg32 mitophagy receptor is a type 2 mitochondrial outer membrane protein which is found on all mitochondria (26, 27), but undergoes post-translational modifications which activate it, presumably on specific mitochondria which are thus marked for degradation (28–30).

An important question is whether the representation of matrix proteins in mitochondria destined for degradation is identical to the average representation in the general mitochondrial network. We previously determined that in *Saccharomyces cerevisiae*, different mitochondrial matrix protein reporters undergo mitophagy at drastically different rates, indicating the existence of a pre-engulfment sorting mechanism (31). We were also able to show that altering mitochondrial dynamics, through deletion of the *DNM1* gene, affected the selectivity that is observed, even though mitochondrial dynamics is not absolutely essential for mitophagy per se, as also confirmed by others (32). These results led us to speculate that mitochondrial dynamics, through repeated mitochondrial fission and fusion cycles, can distill defective components from the network, while sparing other molecular species (31, 33). However, while fission and fusion can ‘shake up’ the network by mixing components, a distillation process would also require some kind of physical segregation principle. A hint into the nature of the putative segregation principle could be related to the function of the mitochondrial phosphatase Aup1 (also known as Ptc6). Aup1 was originally identified by virtue of a synthetic genetic interaction with the Atg1 protein kinase (34). Loss of this gene leads to defects in mitophagy when assayed by some methods, but not by other methods (34, 35). The finding that mitophagy could be protein-specific at the intra-mitochondrial level suggests the possibility that Aup1 could be involved in generating and maintaining mitophagic selectivity, as this would explain the different experimental outcomes obtained using different reporters and conditions. In the present work, we demonstrate that perturbations of mitochondrial protein phosphorylation caused by mutating *AUP1* and additional genes encoding mitochondrial kinases, affect the selectivity of mitophagy. We also show that point mutations at specific phosphosites in a known mitochondrial matrix phosphoprotein, Mdh1, affect the mitophagic efficiency of the protein, and that mitophagic efficiency of Mdh1-GFP is determined downstream of Aup1 function. Thus, our data indicate that targeting of individual protein molecules for mitophagy depends on specific structural determinants and can be further regulated by protein phosphorylation.

## Results

### A role for mitochondrial phosphatases and kinases in regulating the intra-mitochondrial selectivity of stationary phase mitophagy

Aup1, a conserved PP2C- type mitochondrial phosphatase, was previously shown to be required for efficient stationary phase mitophagy, using a simplistic assay which follows the levels of Aco1 (one of two mitochondrial aconitase isoforms in yeast) over the time course of stationary phase mitophagy (34). To test the involvement of Aup1 in mitophagy using more up-to-date methods, we followed the effects of deleting *AUP1* in yeast expressing a chimeric Mdh1-GFP fusion protein. As previously shown for other reporter proteins, delivery of the GFP chimera to the vacuole results in degradation of the native Mdh1 moiety while the residual GFP portion is resistant to vacuolar proteases, generating the appearance of free GFP in immunoblots (36). This can then be used as a semi-quantitative readout of the efficiency of mitophagy (Figure 1). We find that Aup1 is required for mitophagic degradation of Mdh1-GFP, and that this phenotype is complemented by expression of the *AUP1* gene from a plasmid (Figure 1A). Complementation depends on the 34 amino terminal amino acid residues of the protein, which are predicted to contain the mitochondrial targeting sequence. Deletion of the *ATG32* gene completely abrogates the appearance of free GFP (Figure 1B), validating that the free GFP appears specifically as a result of mitophagic trafficking, and not due to another autophagic pathway or other degradation mechanisms (this control was repeated for all mitochondrial GFP fusion proteins used in this study, with identical results; see SI appendix Figure S1).

**Figure 1.**
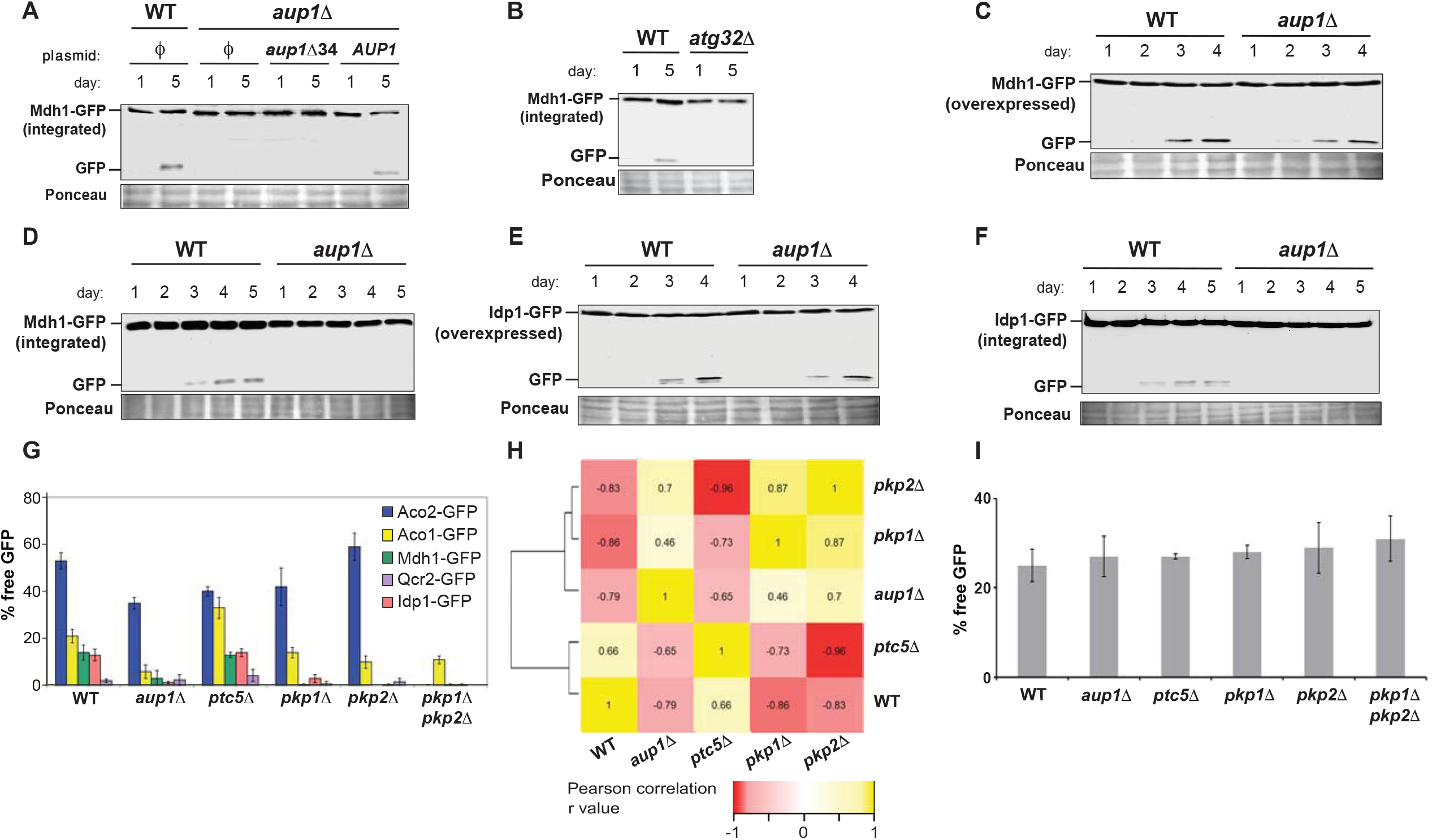
Aup1 and the mitochondrial kinases Pkp1 and Pkp2 play a role in determining protein-level mitophagic selectivity. **(A) deletion of *AUP1* blocks mitophagic trafficking of Mdh1-GFP**. WT (PKY17) and *aup1Δ* (PKY133) cells, harboring different pCU416-derived plasmids as indicated, were grown on SL medium for the indicated times, as described in SI appendix “Supplementary Materials and Methods”. The cells were then harvested and the corresponding protein extracts (20 μg/lane) were subjected to SDS-PAGE and immunoblotted using anti-GFP antibody. Note that truncation of the putative mitochondrial targeting sequence (*aup1Δ34*) of Aup1 prevents the rescue. **(B) Control demonstrating that mitophagic trafficking of integrated Mdh1-GFP is absolutely dependent on the Atg32 receptor protein**. PKY17 cells (WT) and PKY334 cells (*atg32Δ*) were grown and analyzed as in **(A). Panels C-F; Overexpression of reporter proteins suppresses the requirement for Aup1**. WT (HAY75) and *aup1Δ* (HAY809) cells overexpressing Mdh1-GFP **(C)** or Idp1-GFP **(E)** from a plasmid-borne *CUP1* promoter were grown on SL medium for the indicated times, as described in SI appendix “Supplementary Materials and Methods”. The cells were then harvested and the corresponding protein extracts were immunoblotted with anti-GFP antibody. **(D)** and **(F)**; WT and *aup1Δ* cells expressing genome-integrated Mdh1-GFP **(D)** or Idp1-GFP **(F)** from their respective endogenous promoters were grown on SL medium for the indicated times, as described in SI appendix “Supplementary Materials and Methods”. The cells were then harvested and the corresponding protein extracts were immunoblotted with anti-GFP antibody. All blots are representative of at least 3 independent biological replicates. **(G) Mitochondrial kinases and phosphatases regulate the selectivity of mitophagy**. A 5×6 matrix of WT (HAY75), *aup1Δ* (HAY809), *pt5Δ* (PKY37), *pkp1Δ* (PKY61), *pkp2Δ* (PKY70) and pkp1*Δ pkp2Δ* (PKY629) cells, each expressing integrated Aco1-GFP, Qcr2-GFP, Aco2-GFP, Mdh1-GFP or Idp1-GFP from their respective native promoters was assayed for release of free GFP by immunoblotting after a 5 day incubation in SL medium as described in SI appendix “Supplementary Materials and Methods”. Free GFP was quantified by densitometry and normalized as percent of the total GFP signal (free GFP + chimera) in the respective lanes. For specific sample blots which were used in generating these graphs, see SI appendix Figures S2-6. Bars denote standard deviation (n=3, 2-way ANOVA, P=1×10^−17^). For statistical significance of individual pairwise comparisons, see SI appendix Table I. **(H)** Pearson correlation coefficients and clustering analysis for the normalized selectivity vectors defined by each of the single mutant genotypes tested in **(G)** (see Materials and Methods for details). (I) **Deletion of *AUP1*, *PTC5*, *PKP1*, or *PKP2* as well as the *pkp1Δ pkp2Δ* double deletion have no effect on the nonselective mitophagy of an artificial reporter protein**. Cells (HAY72, HAY809, PKY37, PKY61, PKY70 and PKY629) expressing mtDHFR-GFP from plasmid pPKB115 were grown on synthetic lactate medium and mitophagic trafficking of the reporter was determined using % free GFP, as described above. For sample blot, see SI appendix Figure S7.

The defect in mitophagic trafficking of Mdh1 in *aup1Δ* cells can be overcome by overexpressing Mdh1-GFP from a plasmid (Figure 1C-D, and see also SI appendix Figure S12 for a comparison of reporter protein levels in endogenously expressing vs Mdh1-GFP overexpressing strains), suggesting the existence of a saturable selectivity mechanism. This saturation is not a specific property of Mdh1, as the same observation was made when using an Idp1-GFP fusion protein as a reporter (Figure 1E-F). In addition, we tested the effects of deleting Aup1 on the mitophagic trafficking of Aco1, Aco2, and Qcr2, which also localize to the mitochondrial matrix. We find that while Aco1, Mdh1 and Idp1 are affected by knocking out *AUP1*, Aco2 and Qcr2 are less significantly impacted by this mutation (Figure 1G). Yeast mitochondria are known to contain at least two kinases (Pkp1 and Pkp2) and at least two phosphatases (Ptc5 and Aup1). To determine whether the role of Aup1 reflects a broader involvement of protein phosphorylation in determining selectivity, we tested the same 5 reporter proteins for their ability to undergo mitophagy in wild-type, *aup1Δ*, *ptc5Δ*, *pkp1Δ*, and *pkp2Δ* cells (Figure 1G). As previously shown, WT cells exhibit different efficiencies of generation of free GFP for different mitochondrial reporters, under mitophagy-inducing conditions. We then compared the mitophagic selectivity of the different genotypes by calculating pairwise Pearson correlation coefficients, followed by a clustering analysis of the respective data vectors representing each genotype (Figure 1H). We find that, while Ptc5 does not play an appreciable role in regulating mitophagic selectivity (at least with respect to the reporters assayed here), the deletion of either *PKP1* or *PKP2* led to selectivity profiles that were similar to those observed for the deletion of *AUP1* (Figure 1G). This result is consistent with mitochondrial protein phosphorylation playing a general role in determining the intra-mitochondrial selectivity of mitophagic trafficking, not restricted to Aup1. In fact, the double deletion mutant *pkp1Δ pkp2Δ* effectively shuts down mitophagy for most of the endogenous reporters we have tested here (Figure 1G), except for Aco1-GFP, which is inhibited by about 40% in this genetic background. Consistent with this observation, the double mutant exhibited enhanced loss of viability in long incubations on lactate-based medium relative to wild-type (SI appendix Figure S13), reminiscent of data previously published for *aup1Δ* cells (34). In addition, we could show that all of the genotypes tested here showed normal mitophagic trafficking of a totally artificial chimeric mitophagy reporter (mtDHFR-GFP, utilizing mouse DHFR) implying that matrix protein phosphorylation affects the selectivity of mitophagy, as opposed to the process itself, and that the effects of the kinase and phosphatase deletions do not arise from indirect physiological effects which prevent mitophagy or autophagy in general (Figure 1I).

### Mdh1, a mitochondrial matrix protein, is phosphorylated on serine 196, threonine 199 and serine 240 during growth on lactate under mitophagy-inducing conditions

Many mitochondrial matrix proteins are known to undergo phosphorylation (37), although the precise role of these modifications is generally unclear. The apparent involvement of mitochondrial kinases and phosphatases in regulating mitophagic specificity suggests that, at least in some cases, protein phosphorylation affects mitophagic fate. We therefore tested whether Mdh1-GFP, a reporter which shows a high sensitivity to deletion of Aup1, is indeed phosphorylated under our working conditions. Mdh1 contains 5 known phosphosites, at positions 59, 177, 196, 199, and 240 (37–39). As shown in Figure 2, we found by mass spectrometry (MS) analysis of denatured protein extracts that S196, T199 and S240 in Mdh1 are phosphorylated, under these conditions.

**Figure 2.**
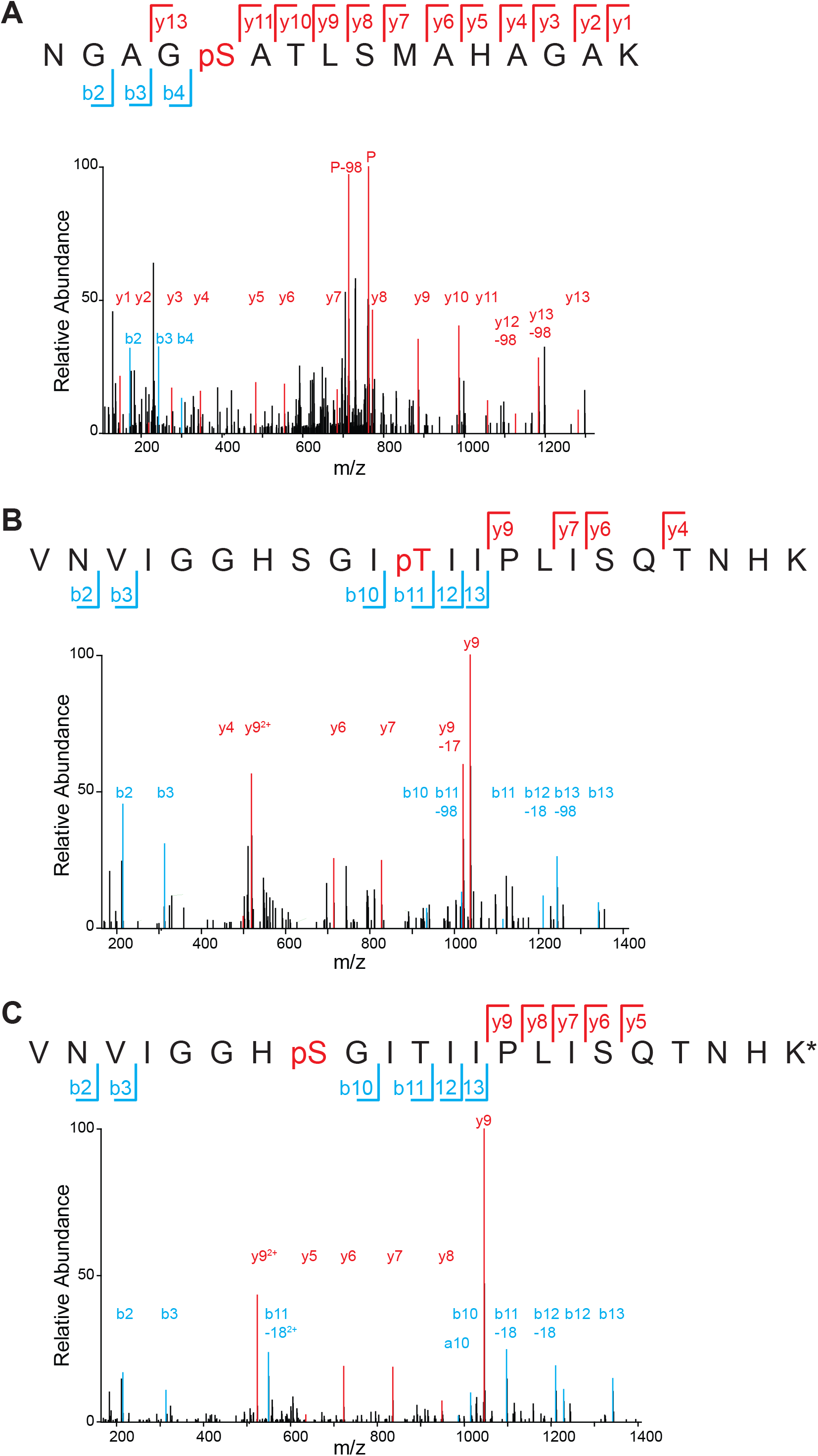
Mdh1-GFP is phosphorylated under mitophagy-inducing conditions. Extracts from TVY1 cells grown in SL medium for 1 day were digested, phosphopeptide-enriched and analyzed by LC-MS/MS for the detection of Mdh1-derived phosphopeptides. **(A**) Serine 240, **(B)** Thr199 and **(C)** Ser196 of Mdh1-GFP are phosphorylated during growth in SL.

### Structural determinants on Mdh1 regulate mitophagic efficiency ‘in cis’

If Mdh1 phosphorylation plays a role in determining mitophagic selectivity, we should expect that mutagenesis of phosphosites on the protein would affect its rate of mitophagy. We carried out site-directed mutagenesis of the sites identified by us (Figure 2), as well as additional reported Mdh1 phosphosites, to the respective alanine and aspartate variants. As shown in SI appendix Figure S11, mutation at these sites did not affect complementation of the growth defect of *mdh1Δ* cells on lactate medium. We find that the T59A, S196A and T199A mutants were unable to undergo mitophagy, while the S240A variant reproducibly showed increased mitophagic efficiency relative to wild-type (Figure 3A). The inability of the T59A, S196A and T199A mutants to undergo mitophagy does not reflect a general block of mitophagy in these cells: a co-expressed mtRFP showed clear induction of mitophagy, as judged by the appearance of red fluorescence in the vacuole, while the T59A and T199A variants maintained a mostly mitochondrial localization (Figure 3B). As an independent verification, we also tested the effect of expressing the T199A variant on the mitophagic trafficking of a co-expressed Qcr2-RFP construct (SI appendix Figure S10): while the T199A mutant is reaching the vacuole less efficiently relative to WT, its expression does not significantly affect mitophagy of the Qcr2-RFP construct in the same cell. The fact that T199A is defective in reaching the vacuole (Figure 3B) makes it unlikely that the point mutants are delivered to the vacuole, but for some reason are recalcitrant to degradation by vacuolar proteases. To further rule out the possibility that differences in sensitivity to vacuolar proteases underlie the difference in release of free GFP from Mdh1-GFP between WT and the T199A mutant, we deleted the mitochondrial targeting sequence in these constructs. The resultant Mdh1-GFP molecules are cytoplasmic, and can be directly targeted to the vacuole by nitrogen-starvation-induced macroautophagy, thus completely bypassing mitochondria (40). If the difference observed between WT Mdh1-GFP and the T199A mutant is due to differential sensitivity to vacuolar proteases, then we expect it to persist in constructs lacking the MTS under general macroautophagy-inducing conditions, as the path taken en route to the vacuole should not affect the sensitivity to proteolysis within the vacuole. However, in the experiment shown in Figure 3C, we can see that both MTSΔ constructs generate very similar amounts of free GFP upon nitrogen starvation, in stark contrast with the respective mitochondrially-targeted constructs. The free GFP which is formed from the MTSΔ constructs is indeed due to direct macroautophagic transport from the cytosol to the vacuole, as it is independent of Atg32, but totally dependent on Atg1. Finally, the point mutations used in Figure 3A had no effect on cell growth (SI appendix Figure S11), indicating that these proteins are enzymatically active. Thus, we can conclude that discrete changes to a protein’s structure can have a clear effect on its mitophagic targeting without affecting mitophagic targeting of other mitochondrial proteins.

**Figure 3.**
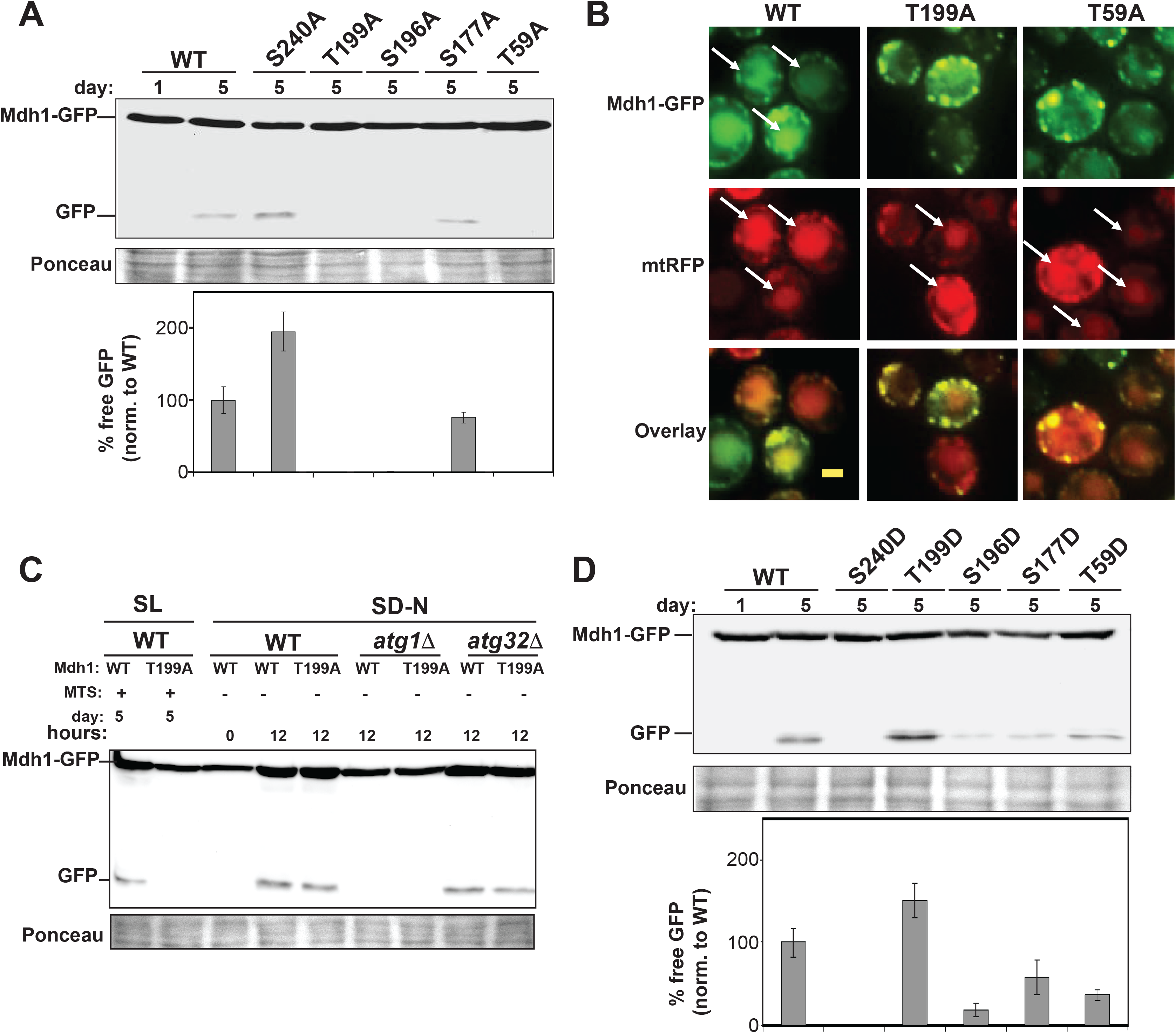
Serine/threonine to alanine mutagenesis of known phosphosites on Mdh1 affects the selective mitophagy of Mdh1-GFP without blocking general mitophagy. **(A)** PKY365 cells harboring centromeric plasmids expressing the different Mdh1-GFP phosphosite alanine mutants from the endogenous *MDH1* promoter were incubated in SL medium for 1 or 5 days as indicated, and protein extracts were prepared. Equal amounts of protein (20 μg) were subjected to SDS-PAGE and immunoblotting with anti-GFP antibody. For quantification, the % free GFP was normalized to WT, for each variant. Error bars indicate S.E. (n=3). **(B) Point mutations in Mdh1-GFP prevent mitophagic trafficking of the specific reporter, but do not affect mitophagic trafficking of a co-expressed mtRFP reporter**. Cells (PKY365) expressing WT Mdh1-GFP and the indicated variants from the native *MDH1* promoter as well as a mitochondrially targeted mtRFP were incubated in SL medium for 5 days and imaged by fluorescence microscopy. WT Mdh1-GFP shows mitophagic targeting (white arrows) by day 5, which co-localizes with the mtRFP signal in these cells. In contrast, the mutants show defective routing to the vacuole, while the RFP signal in the vacuole is not affected (white arrows). Scale bar = 1 μm. **(C) The difference between point mutants of Mdh1 cannot be explained by differential sensitivity to vacuolar proteases, and depends on mitochondrial targeting**. WT (HAY75), *atg32Δ* (HAY1239), and *atg1Δ* (HAY339) cells expressing truncated versions of Mdh1 which lack the mitochondrial targeting sequence were subjected to nitrogen starvation for 12 h to induce macroautophagy, and protein extracts were analyzed by immunoblotting. The two left-most lanes are control cells expressing full length WT and T199A versions of Mdh1, which were subjected to the standard 5 day stationary phase mitophagy protocol, for comparison (N=3). **(D) Serine/threonine to aspartate mutagenesis of T199 and S240 on Mdh1 leads to reciprocal effects on Mdh1-GFP mitophagic trafficking, relative to the respective alanine mutations**. PKY365 cells transformed with vectors expressing Mdh1-GFP and the indicated variants from the endogenous *MDH1* promoter, were incubated in SL medium for 1 or 5 days, and protein extracts were prepared. Equal amounts of protein (20 μg) were subjected to SDS-PAGE and immunoblotting with anti-GFP antibody. For quantification, the % free GFP was normalized to WT for each variant. Error bars indicate s.d, analysis of variation of three independent experiments (N=3).

Consistent with these findings, some of the corresponding aspartate mutations showed a reciprocal effect. Thus, mitophagic targeting of the S240D mutant is strongly attenuated relative to the wild type, and the mitophagic targeting of the T199D mutant is increased, relative to wild-type (see Figure 3D).

### Time and Aup1-dependent changes in the phosphorylation state of Mdh1-GFP occur during mitophagy

To further understand the role of matrix phosphorylation in mitophagic selectivity, we analyzed time- and genotype- dependent changes in the phosphorylation state of Mdh1-GFP during mitophagy. Native immunoprecipitation of Mdh1-GFP from cell extracts, followed by immunoblotting with anti-phosphoserine/phosphothreonine antibody (anti pSer/pThr) identifies a λ-phosphatase sensitive signal which increases in WT cells over the incubation period, but decreases over time in the isogenic *aup1Δ* strain, and is undetectable by day 2-3 of the incubation, when mitophagy is normally induced (Figure 4A). We also managed to quantify peptides containing phospho-Ser196 and phospho-Thr199 by SILAC-based MS analysis, comparing WT to *aup1Δ* cells. At day 2, phosphoSer196 appears to be stable, whereas phosphoThr199 is significantly less abundant in *aup1Δ* cells relative to *AUP1* expressing cells (Figure 4B). Thus, we conclude that the reduced phosphorylation of Thr199 contributes to the decrease of phosphoSer/Thr signal observed by western blot analysis of anti-GFP immunoprecipitates in *aup1Δ* cells.

**Figure 4.**
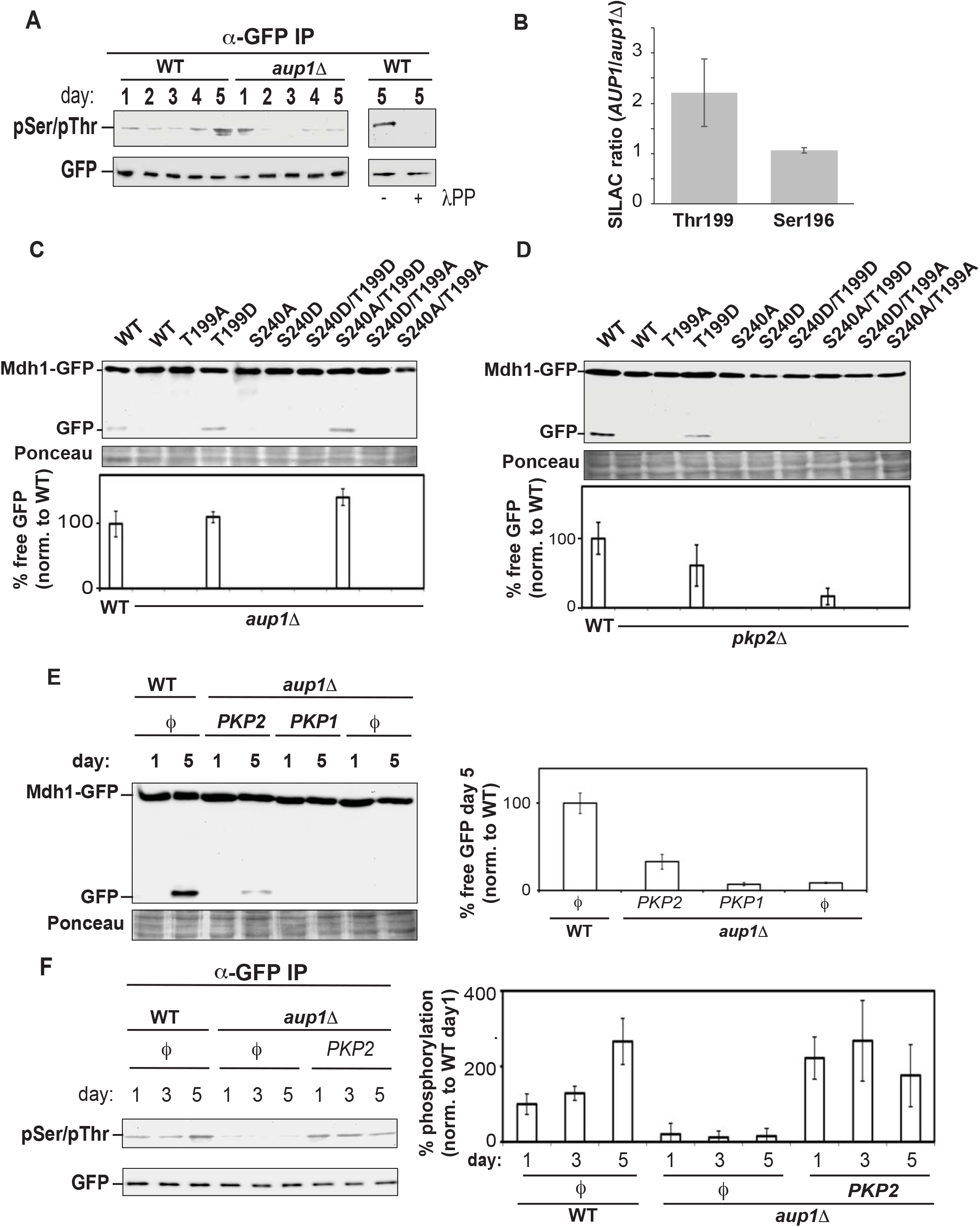
Aup1 functions upstream of Pkp2, to generate dynamic changes in Mdh1 phosphorylation which are required for mitophagic targeting. **(A) Anti-phosphoamino acid immunoblotting of anti-GFP immunoprecipitates demonstrate the effects of *AUP1* deletion on Mdh1-GFP phosphorylation**. PKY395 (*aup1Δ*) and isogenic control TVY1 cells, both expressing Mdh1-GFP, were grown in SL medium. At the indicated time points, protein extracts were generated under native conditions and immunoprecipitated with anti-GFP antibodies as detailed in “Materials and Methods”. The immunoprecipitates were analyzed by immunoblotting with anti-phosphoamino acid antibody (top) and anti-GFP antibody (bottom). Where noted (right panel), anti-GFP Immunoprecipitates were treated with lambda phosphatase for 2 h. All data represent a minimum of 3 independent biological replicates. **(B) SILAC analysis of relative phosphorylation levels at position S196 and T199 of Mdh1-GFP on day 2 of incubation in SL medium**. Samples were collected from *AUP1* (strain PKY520, heavy) and *aup1Δ* (strain PKY661, light) cells as described in “Methods”, and anti-GFP immunoprecipitates were analyzed by LC-MS/MS to determine the ratio of the indicated phosphopeptides in WT versus *aup1Δ* cells. **(C-D) Expression of the ‘hypermitophagic’ *MDH1* mutant T199D can suppress the *aup1Δ* and *pkp2 Δ* phenotypes**. *C*ells (PKY365) expressing WT Mdh1-GFP, as well as **(C)** *aup1Δ* (PKY456) or *pkp2Δ* (PKY488) cells expressing WT Mdh1-GFP and the indicated Mdh1 variants (all constructs were expressed from the endogenous *MDH1* promoter) were incubated in SL medium for 5 days and protein extracts were prepared. Equal amounts of protein (20 μg) were subjected to SDS-PAGE and immunoblotting with anti-GFP antibody. The % free GFP was normalized relative to WT Mdh1-GFP, for each variant. **(E) Overexpression of Pkp2, but not Pkp1, can bypass the *aup1Δ* block in Mdh1 mitophagic trafficking**. Pkp1 and Pkp2 were overexpressed from the *CUP1* promoter. **(F) Overexpression of Pkp2 leads to recovery of Mdh1-GFP phosphorylation in the *aup1Δ* background**. TVY1 (*AUP1* control) and PKY395 (*aup1Δ*) cells, expressing Pkp2 or harboring empty vector (*ϕ*) were grown in SL medium for the indicated times and Mdh1-GFP was immunoprecipitated under native conditions and analyzed by immunoblotting with anti pSer/pThr antibodies. Data is an average of 3 biological replicates; asterisks denote p≤0.05 (single tail t-test, N=3).

### Expression of a hyper-mitophagy variant of Mdh1-GFP can suppress mitophagy defects in *aup1Δ* and *pkp2Δ* cells

If phosphorylation of Mdh1 at T199 is important for mitophagic trafficking of the protein, and if Aup1 directly or indirectly regulates the phosphorylation state at this residue, then we expect the T199D mutant to be able, at least partially, to bypass the phenotype of the *aup1Δ* or *pkp2Δ* mutations. As shown in Figure 4C and 4D, the T199D mutation shows reproducible suppression of both the *aup1Δ* and *pkp2Δ* mutants. However, neither of the position 240 mutants was able to bypass the *aup1Δ* mutation. These results clearly indicate that modification of Mdh1 at position 199 can regulate mitophagic trafficking of Mdh1-GFP downstream of Aup1 and Pkp2. For the effects of expressing these constructs in WT cells, see SI appendix Figure S9.

### Overexpression of Pkp2, but not Pkp1, can bypass the mitophagic defect of Mdh1-GFP in *aup1Δ* cells

Deletion of *AUP1* leads to hypo-phosphorylation of Mdh1-GFP, as judged by anti-phosphoamino acid immunoblot (Figure 4A). This makes it implausible that the effects of Aup1 are due to direct dephosphorylation of Mdh1 by the phosphatase. Alternatively, *AUP1* could regulate Pkp1 and/or Pkp2 which will, in turn, regulate the phosphorylation state of Mdh1. To test this second possibility, we overexpressed either Pkp1 or Pkp2 in *aup1Δ* cells, and assayed delivery of Mdh1-GFP to the vacuole under mitophagy inducing conditions through release of free GFP. As shown in Figure 4E, the overexpression of Pkp2, but not Pkp1, allowed free GFP release from Mdh1-GFP in *aup1Δ* cells. This result is consistent with Pkp2 functioning downstream of Aup1, in regulating the mitophagic clearance of Mdh1-GFP. In further support of this interpretation, overexpression of Pkp2 rescues the hypophosphorylation of Mdh1-GFP in *aup1Δ* cells (Figure 4F): While *aup1Δ* cells harboring empty vector show hypophosphorylation of Mdh1-GFP, *aup1Δ* cells overexpressing Pkp2 show a clear recovery of the anti-phosphoamino acid immunoblot signal. This result strongly indicates that Aup1 regulates Pkp2 activity, which in turn regulates the phosphorylation state of Mdh1-GFP.

### Deletion of *AUP1* leads to changes in the protein interaction network of Mdh1-GFP

To probe the mechanism by which changes in phosphorylation pattern affect mitophagic trafficking of Mdh1-GFP, we compared its interactomes in WT and *aup1Δ* cells by quantitative proteomics. Anti-GFP immunoprecipitations from SILAC-labeled WT and *aup1Δ* cells grown on lactate were performed in the presence or absence of Mdh1-GFP expression, followed by MS analysis. The raw MS data are shown in supplemental table V. As shown in Figure 5A, most of the significant interactors of Mdh1-GFP appear above the diagonal, indicating that generally, Mdh1-GFP interacts more strongly with these proteins in the *aup1Δ* background, at least on day 1. We further validated individual interactions by tagging putative interaction partners with the HA epitope, and assaying native co-immunoprecipitation with Mdh1-GFP. The individual interactions tested included the previously known interactor Cit1-HA, the newly identified interactor Idh1-HA, and Mic60-HA, a known Mdh1 interactor which did not come up in our SILAC screen (41, 42). In all three cases, we were able to recapitulate the general interactome data, as all interactions (in day 1 samples) were stronger in the mutant, relative to WT (Figure 5B-D). The Cit1-Mdh1 interaction was far stronger on day 1 in the *aup1Δ* mutant than in WT, and increased on day 4 in the mutant, relative to day 1 (Figure 5B). Idh1-HA also showed stronger interaction with Mdh1-GFP in the day 1 samples (Figure 5C) but this reversed on day 4. Finally, the Mic60 interaction showed increased interaction in the mutant, but weakened considerably between days 1 and 4 of the incubation (Figure 5D) such that the interaction in WT cells became undetectable in the day 4 sample. Thus, deletion of *AUP1* appears to have dynamic and divergent effects on the Mdh1 interactome, which may underlie its effects on mitophagic targeting of the protein.

**Figure 5.**
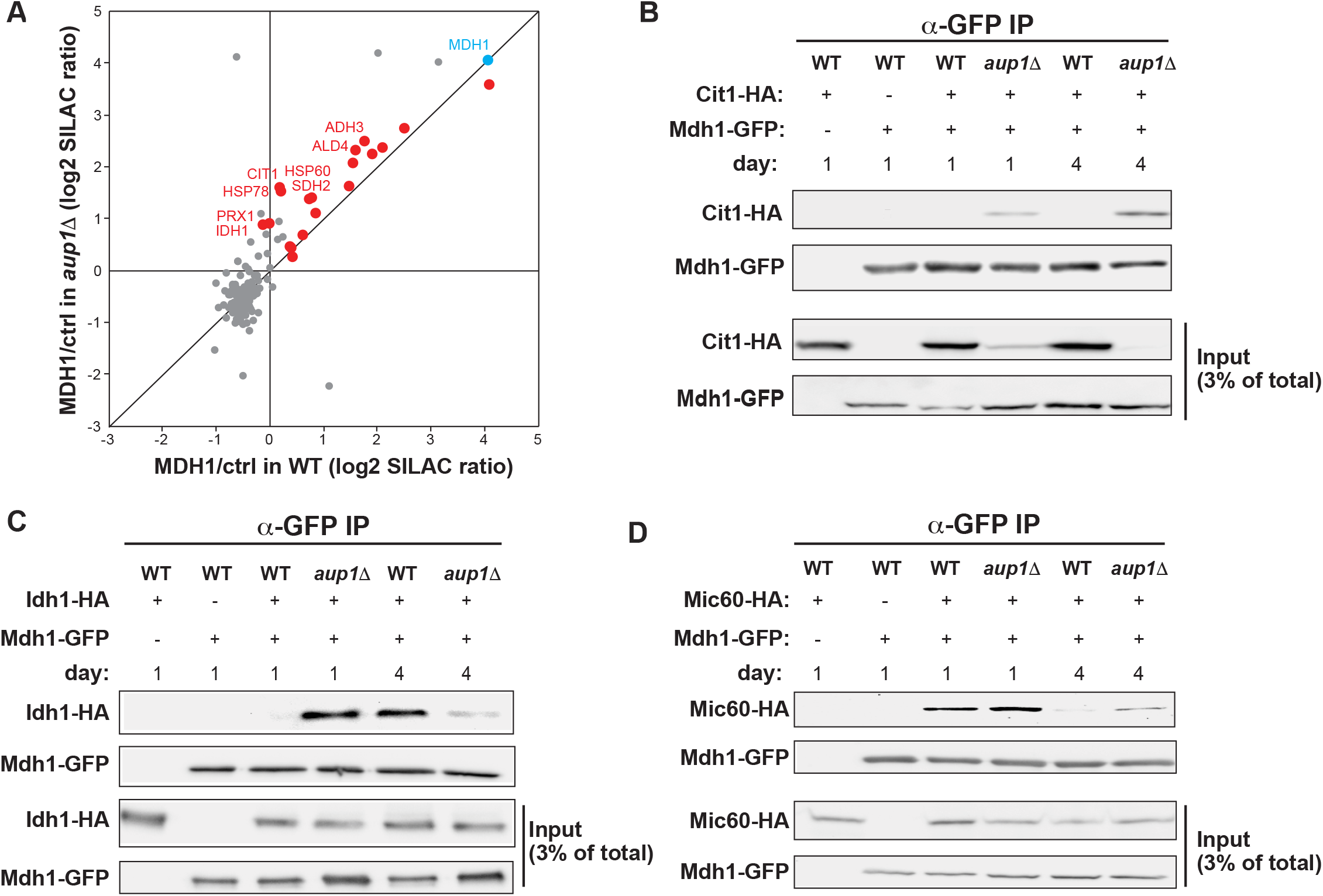
Loss of Aup1 affects protein-protein interactions of Mdh1. **(A)** Interactome analysis of Mdh1-GFP in WT and *aup1Δ* cells using SILAC. Heavy isotope labeled Mdh1-GFP was immunoisolated from PKY392 (*AUP1* WT control) and PKY415 (*aup1Δ*) cells after 1 day of growth in synthetic lactate medium. Prior to protein extraction, the cells were mixed with an equal amounts of isogenic untagged cells (light isotope labeling) as described in “Materials and Methods”. Specifically interacting proteins were identified by comparing MDH1-GFP affinity purifications to cells not expressing GFP-tagged MDH1 (ctrl; n=2). Significantly interacting mitochondrial matrix proteins are highlighted in red (p<0.05, outlier test). All other identified proteins are marked in grey. Significantly enriched matrix proteins that change their interaction by at least 1.5-fold depending on the presence of AUP1 are annotated. **(B-D)** Wild-type and *aup1Δ* cells (PKY826, PKY827, PKY824, PKY825, PKY1032, PKY1040) expressing either Cit1-HA, Idh1-HA, or Mic60-HA from the respective endogenous promoters were grown for 1 or 4 days in synthetic lactate medium, and native extracts (prepared as detailed in “Materials and Methods”) were immunoprecipitated with anti-GFP antibodies. The immunoprecipitates were analyzed by immunoblotting with anti-HA antibodies. **(B)** loss of Aup1 affects the interaction between Mdh1 and Cit1. **(C)** loss of Aup1 affects the interaction between Mdh1 and Idh1. **(D)** loss of Aup1 affects the interaction between Mdh1 and Mi60. Data are representative of at least 3 independent biological replicates.

### Deletion of *AUP1*, and point mutations which abrogate mitophagic trafficking of Mdh1, alter the distribution of the Mdh1-GFP reporter protein within the mitochondrial network

We previously published that GFP chimeras of protein species which undergo inefficient mitophagy appear to be segregated within the network, relative to a generic mitochondrial RFP reporter (31). We therefore wanted to test whether the molecular perturbations which were found to affect mitophagic trafficking in the present study, also affect the distribution of the Mdh1-GFP reporter protein within the mitochondrial network. As shown in Figure 6A, there is a clear difference in the distribution of Mdh1-GFP between WT cells and *aup1Δ* cells at day 2 of the incubation, with *aup1Δ* cells exhibiting apparent clumping of the GFP signal to discrete dots within the mitochondrial network as defined by the mtRFP signal. Consistent with this, Figure 6B illustrates that wild-type Mdh1-GFP, and a point mutant which does not reduce mitophagic trafficking, (S240A), show a statistically significant increase in the signal overlap of the red and the green channels between day 1 and day 2 of the incubation, as determined using the JACoP plugin (43). In contrast, we observed that mutants which do show a significant block in mitophagic trafficking display a statistically significant decrease in signal overlap (relative to mtRFP) from day 1 to day 2, similar to that observed in the *aup1Δ* strains. It is important to note that none of the mutations tested had any effect on the general fractionation properties of Mdh1-GFP (see SI appendix Figure S8) making it unlikely that they affect targeting of the protein into mitochondria. Rather, the data is consistent with some degree of segregation of these mutants, relative to the overall available space in the mitochondrial matrix as defined by the mtRFP fluorescence signal. In addition, we were able to directly follow segregation of Mdh1-RFP and Qcr2-GFP within the same cells (SI appendix Figure S14), confirming that segregation of these endogenously expressed reporter proteins correlates with their different mitophagic efficiencies.

**Figure 6.**
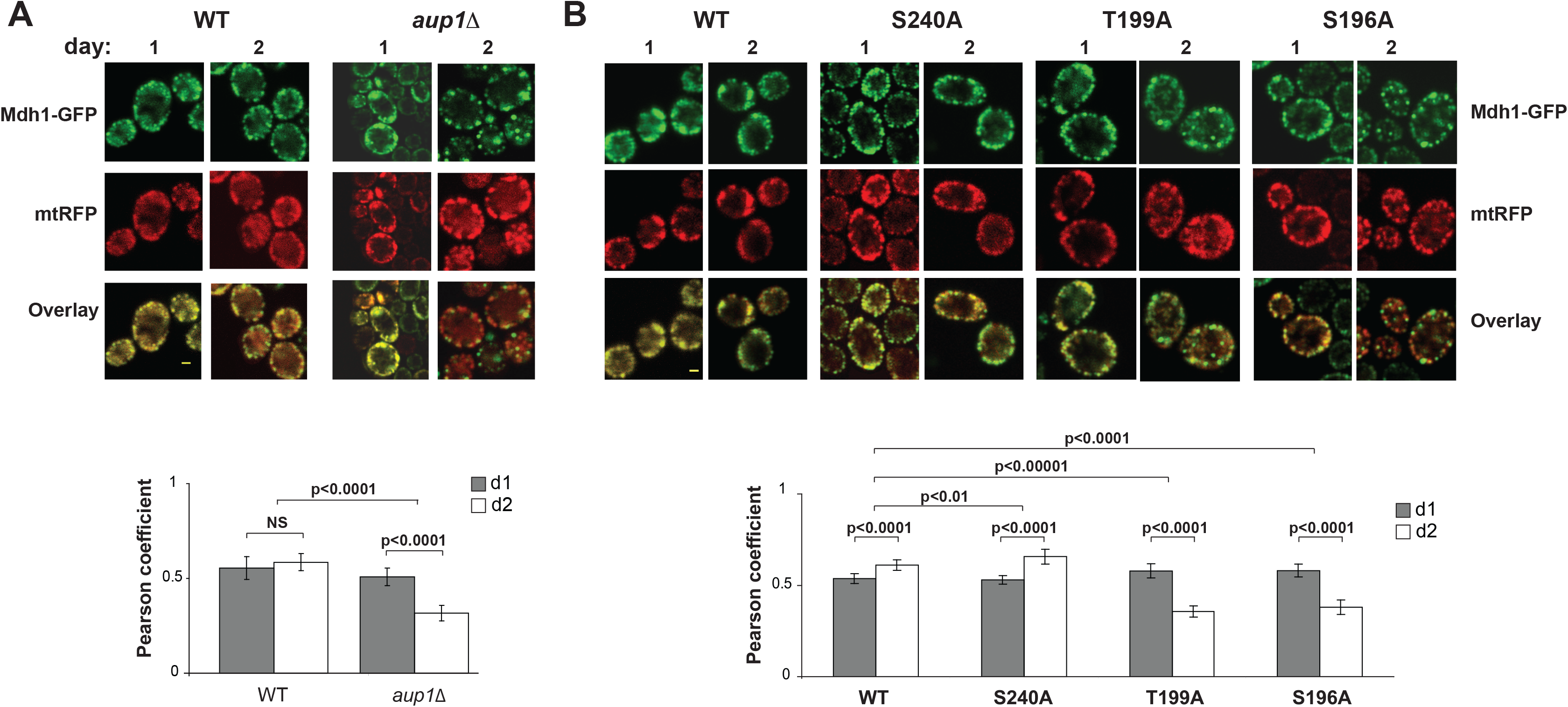
Phosphorylation state affects the matrix distribution of Mdh1-GFP. **(A) Deletion of *AUP1* affects distribution of Mdh1-GFP within the mitochondrial matrix**. Cells (*mdh1Δ*) co-expressing Mdh1-GFP from the endogenous promoter and a generic mtRFP were imaged on days 1 and 2 of the incubation (to avoid interference from vacuolar signal, top panels) and the overlap between the red and green channels was quantified as described in ‘Materials and Methods’. Graph (bottom) shows quantification of the overlap between the two channels and statistical significance (NS, not significant). **(B) The T199A and S196A mutations also increase the segregation of Mdh1-GFP, relative to a matrix mtRFP**. Cells (*mdh1Δ*) expressing the indicated Mdh1-GFP species and a generic mtRFP were imaged and analyzed as in **A**. Results are representative of two biological replicates. Scale bar =1 μm.

## Discussion

The finding that within an individual protein species, differentially modified forms of individual protein species can have different mitophagic fates constitutes an important advance in our mechanistic understanding of protein-level mitophagic selectivity. It suggests that specific ‘degron-like elements’ on mitochondrial matrix proteins are regulating selectivity, and this now allows further analysis of the molecular mechanism. While individual mitochondrial phosphatases and kinases each affect a subset of mitochondrial proteins, sometimes incompletely, we found that knockout of both mitochondrial kinases (Pkp1 and Pkp2) completely blocked mitophagic trafficking of nearly all the reporters tested in this study. Therefore, we can conclude that matrix protein phosphorylation is a widespread determinant of mitophagic selectivity.

We further demonstrate that a novel protein phosphorylation cascade plays a role in the regulation of selectivity. The Aup1 phosphatase functions upstream of the Pkp2 kinase in this cascade, as (1) the Mdh1^T199D^ mutant was able to partially bypass the specific requirements for both Aup1 and Pkp2 (Figure 4, panels C and D); (2) it is unlikely that direct dephosphorylation of threonine 199 by Aup1 is responsible, since the phosphomimetic mutations point to the phosphorylated form of T199 as being ‘mitophagy competent’ (Figures 3A, 3D, 4C, and 4D). It is much more likely that Aup1 is required, directly or indirectly, for the activation of a kinase such as Pkp2. Indeed, we found that overexpression of Pkp2, but not Pkp1, can suppress both the Mdh1 mitophagy defect (Figure 4E) and the hypophosphorylation of Mdh1 (Fig. 4F), which are observed in *aup1Δ* cells. This strongly suggests that Aup1 activity is required for activation of Pkp2, and that Pkp2 directly phosphorylates Mdh1 at position 199. The fact that the S240D mutation is able to prevent rescue of the *aup1Δ* phenotype by T199D in the context of the S240D T199D double mutant (Figure 4C) may suggest that the two sites represent a combinatorial output, not a strictly hierarchical one where position 240 affects the phosphorylation state of threonine 199.

The two known yeast mitochondrial kinases (Pkp1 and Pkp2) have been identified as orthologs of mammalian PDKs (pyruvate dehydrogenase kinases), which exist as a family of paralogs with different tissue distribution in metazoans (44). Reports in the literature (45, 46) also link the two phosphatases, Ptc5 and Aup1 (Ptc6) to Pyruvate Dehydrogenase Alpha subunit (Pda1) dephosphorylation in yeast. We were unable to test for a direct role of Pda1 in mitophagy, as the deletion mutant shows severely impaired growth in our liquid SL medium. Nonetheless, our data are not consistent with Pda1 phosphorylation being the determinant of mitophagic selectivity and efficiency in our experiments, for the following reasons. First of all, yeast Pda1 has only one verified phosphosite which is regulated by these enzymes: serine 313. Thus, if all of these mitochondrial kinases and phosphatases in yeast converge exclusively on the regulation of Pda1, then we expect Aup1 and Ptc5 to have opposite phenotypes relative to Pkp1 and Pkp2. However, our results are inconsistent with this prediction, and indicate that Aup1 functions to regulate and activate Pkp2 in our readouts. In addition, the apparent lack of involvement of Ptc5 and the non-redundant role of Aup1 in the phenotypes observed also argue against an interpretation which converges on Pda1 phosphorylation. Finally, one should note that there are hundreds of documented phosphorylation sites in yeast mitochondrial proteins (37, 47) and it is therefore unlikely that the only role of Aup1, Ptc5, Pkp1 and Pkp2 is to regulate phosphorylation of serine 313 on Pda1. While Pkp1 and Pkp2 are separately and non-redundantly required for mitophagic trafficking of Mdh1-GFP (see Figure 1G), only Pkp2 overexpression can partially bypass the *aup1Δ* block (Figure 4E). This either implies that Pkp1 functions in a separate Aup1-independent pathway, or that Pkp1 is under additional regulation such that its overexpression per se does not lead to sufficiently high activity levels.

Yeast cells are not the only system in which protein-level specificity was observed during mitophagy. Hämäläinen *et al* (48) showed that in differentiation of MELAS patient-derived iPSCs into neurons, mitochondrial respiratory chain (RC) complex I (CI) underwent a selective mitophagic degradation process which spared other RC components. In addition, they could show that the remaining CI in these cells was segregated into distinct patches within the mitochondrial network, while other RC components were evenly distributed. Their results demonstrate that intra-mitochondrial selectivity of mitophagic degradation also occurs in mammalian cells, and is not restricted to yeast. Interestingly, phosphorylation of complex I components has been reported in mammalian cells (49). In cultured mammalian cells, Burman et al demonstrated selective mitophagic clearance of a mutant ornithine transcarbamoylase (50). Finally, Vincow et al (51, 52), using *Drosophila*, found that mitophagy is protein selective in a whole animal system.

The combined loss of Pkp1and Pkp2 has a reproducible effect on yeast survival in mitophagy-inducing conditions (SI appendix Figure 13) which is consistent with previously published data for *aup1Δ* cells (34). The precise mechanism by which defects in mitophagic selectivity affect cell survival will be determined in future studies, and may hypothetically involve a toxic accumulation of defective proteins or other molecular species.

How can we envision the effects of posttranslational matrix protein modification on mitophagic selectivity of individual protein molecules? We previously proposed that phase separation within the mitochondrial matrix, coupled with repeated fission and fusion cycles, would lead to a distillation effect where the “minority” phase would be segregated and restricted to homogeneous mitochondrial compartments (31, 33). If phosphorylation, or any other modification, affects the partitioning of a given protein molecule between these phases, then it would also affect the protein’s segregation tendencies. We then need to postulate that the accumulation of proteostatic load in a given mitochondrial compartment leads to localized activation of mitophagy receptors, leading to selective degradation of mitochondrial compartments enriched in specific protein species. This outlines a model in which protein lesions lead to changes in posttranslational modifications of the damaged molecules, which lead in turn to these proteins being concentrated into discrete compartments that are targeted for mitophagic degradation (Figure 7). Regulation of phase partitioning of proteins by phosphorylation has precedents: the regulation of RNA granule formation in the cytosol (53–55) and milk globule formation (56) are two well-studied examples.

**Figure 7.**
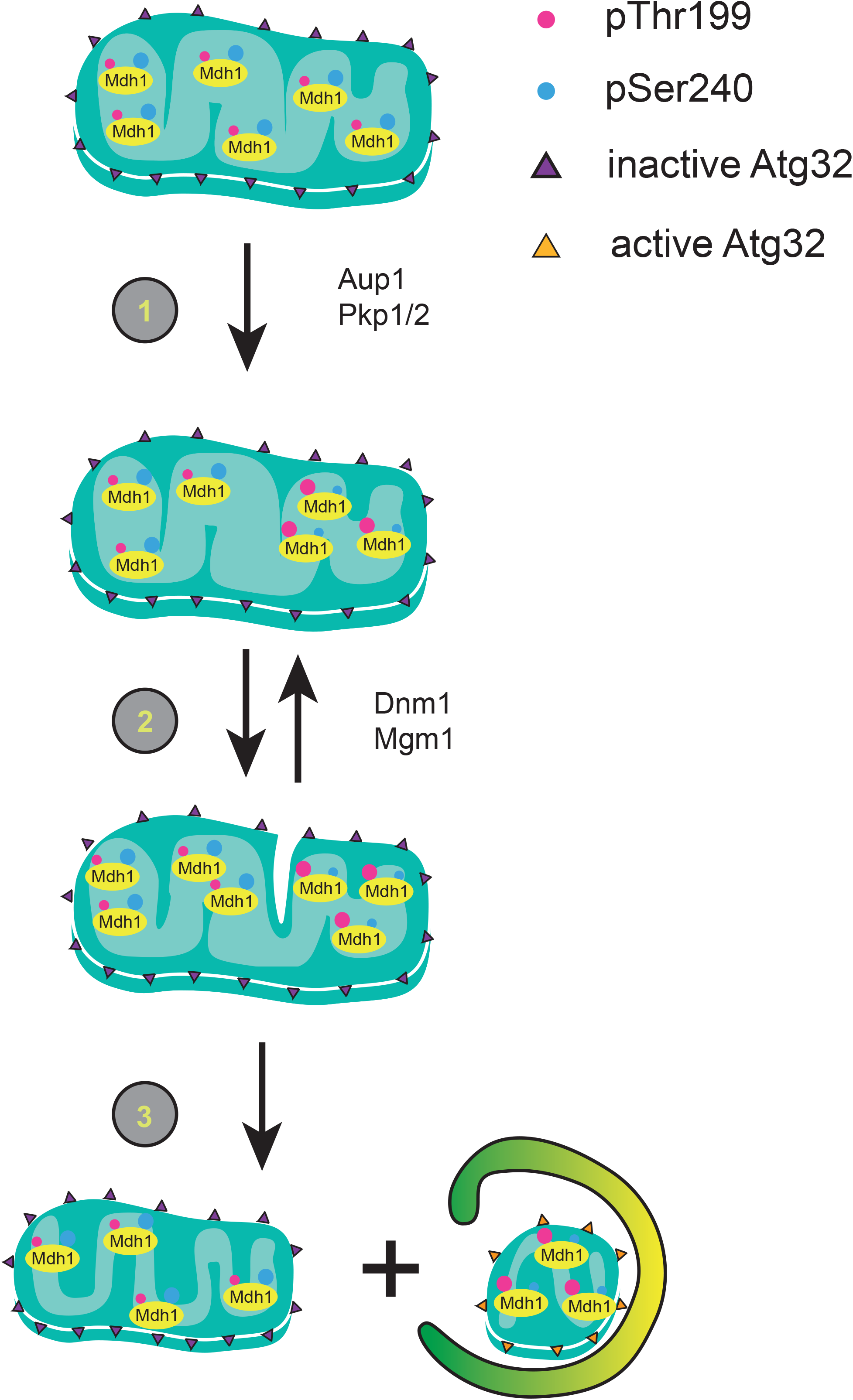
A model for the role of protein phosphorylation in regulating mitophagic selectivity. We suggest that phosphorylation of mitochondrial matrix proteins such as Mdh1 can regulate their tendency to undergo mitophagy, within the matrix. **(1)** Increased phosphorylation of T199 and decreased phosphorylation at S240 bring about a change in protein-protein interactions that leads to segregation of this species. **(2)** multiple fusion and fission events lead to a ‘distillation’ like process that enriches this species in a single mitochondrial compartment, along with other protein species with similar properties. **(3)** A final fission event generates a mitochondrion sufficiently enriched in ‘degradation-bound’ species. We hypothesize that the proteostatic load in this compartment generates a signal which activates the Atg32 receptor, possibly through Yme2-dependent clipping of the Atg32 C-terminus, leading to engulfment of this compartment- with its enclosed protein components- by autophagic sequestering membranes.

Such a model, where fission and fusion play an indirect role in mitophagy, would reconcile a number of apparently contradictory results in the literature. Many publications have claimed that mitochondrial fission is essential for mitophagy (57–60). However, most of these posit a direct mechanistic role, either in generating ‘bite sized’ engulfable fragments and some even imply a molecular interaction between the autophagic machinery and the fission machinery (61). In contrast, a recent publication claimed that mitochondrial fission mediated by Drp1/Dnm1 is completely unnecessary for mitophagy (32), and that the autophagic machinery itself mediates fission during engulfment. A model in which mitochondrial dynamics has no role whatsoever does not explain the reproducible slowdown of mitophagy in *dnm1Δ* cells, which was repeatedly observed in multiple publications from different groups (31, 32, 62). However it is perfectly consistent with a ‘percolation’ or ‘distillation’ role of mitochondrial dynamics, in determining the selectivity and rate of mitophagy, as we have proposed (31, 33).

An alternative hypothesis would be to postulate that specific protein-protein interactions regulate the segregation of proteins into mitophagy-bound compartments. This type of model must address several difficult questions. Are all mitochondrial proteins interacting with a small subset of proteins which function promiscuously to mediate mitophagic selectivity? Surely one cannot have each mitochondrial matrix protein surveilled by a different and unique mitophagy regulator? We currently favor the nonspecific, phase separation model, as it imposes fewer constraints on the system and requires fewer assumptions. However, further studies must be carried out before one of these hypotheses is ruled out.

In summary, our results suggest a potential mechanism for generating mitophagic selectivity at the molecular level, and indicate a role for protein phosphorylation in regulating mitophagic trafficking of individual mitochondrial matrix protein molecules.

## Materials and Methods

### Plasmids and DNA manipulations

The plasmids and oligonucleotides used in this study are listed in SI appendix Tables III and IV, respectively. For individual plasmid construction and site directed mutagenesis, see SI appendix.

### Yeast strains and growth conditions

Yeast strains used in this study are listed in SI appendix Table II. Deletion mutants and epitope-tagged strains were generated by the cassette integration method (63). Strain HAY75 (MATα *leu2-3,112 ura3-52 his3-Δ200 trp1-Δ901 lys2-801 suc2-Δ9*) was used as the WT genetic background in this study. For primer details, see SI appendix table III.

All knockout strains were verified by PCR. Oligonucleotides used in this study are detailed in SI appendix Table IV. For detailed culture growth conditions, please see SI appendix.

### Chemicals and antisera

Chemicals were purchased from Sigma-Aldrich (Rehovot, Israel) unless otherwise stated. Custom oligonucleotides were from Hylabs (Rehovot, Israel). Anti-GFP and anti-HA antibodies were from Santa-Cruz (Dallas, TX). Anti-phospho serine/threonine antibodies were from Abcam (Cambridge, UK). Horseradish peroxidase-conjugated goat anti-rabbit antibodies were from MP Biomedicals (Santa Ana, CA) or Jackson ImmunoResearch (West Grove, PA).

### Fluorescence microscopy

Culture samples (3 μl) were placed on standard microscope slides and viewed using a Nikon Eclipse E600 fluorescence microscope equipped with the appropriate filters and a Photometrics Coolsnap HQ CCD. To achieve statistically significant numbers of cells per viewing field in the pictures, 1 ml cells were collected and centrifuged for 1 min at 3,500 xg. For quantitative analysis of co-localization, ImageJ software with the co-localization Plugin, JACoP (43). Representative fields were analyzed in terms of channel intensity correlation coefficient. For confocal microscopy, cells were placed on standard microscope slides and micrographs were obtained by confocal laser scanning microscopy using a Leica SP8, using a 63x water immersion lens. GFP excitation was carried out using the 488 nm line, and emission was collected between 500-540 nm. For RFP excitation we used the 552 nm line and fluorescence emission was collected between 590-640 nm. GFP and RFP co-localization images were acquired using LasAF software and the images were analyzed by Fiji software. For statistical analysis of channel overlap data, ANOVA analysis was carried out using JMP 12 software. Correlations were compared over mutations and days by two-factor ANOVA. As the mutation X day interaction effect was statistically significant (p<0.0001), the two days were compared for each mutation by contrast t-tests. Pre-planned comparisons to WT were also performed by contrast t-tests.

### Native immunoprecipitation

Cells (20 OD_600_ units) were washed 3x with 1 ml of cold 10 mM PIPES pH 7, 2 mM PMSF. The cell pellet was washed in three volumes of cold lysis buffer (20 mM HEPES pH 7.4, 50 mM NaCl, 1mM EDTA, 10% glycerol, 0.2 mM Na_3_VO_4_, 10 mM NaPO_4_, 10 mM β-glycerophosphate, 10 mM NaF, 10% phosphate cocktail inhibitors, 1 mM PMSF, proteases inhibitors (final concentrations: 5 μg/ml antipain, 1 μg/ml aprotinin, 0.5 μg/ml leupeptin, 0.7 μg/ml pepstatin). The cells were then frozen in liquid nitrogen and stored at −80°C. Extracts were prepared from the frozen pellets using a SPEX Freezer/Mill according to the manufacturer's instruction. The frozen ground yeast powder was thawed at room temperature, and equal amount of lysis buffer plus Triton-X100 was added to a final concentration of 0.5% v/v detergent. Cell debris was removed by centrifugation at 17.000x g for 10 min at 4°C. 20 μl of GFP-Trap bead slurry (Chromotek) was pre-equilibrated in lysis buffer, per the manufacturer's instructions. The cleared lysate was added to the equilibrated beads and incubated for 1 h at 4 °C with constant mixing. The beads were washed five times with 1 ml ice cold wash buffer (20 mM HEPES pH 7.4, 50 mM NaCl, 0.5 mM EDTA, 10% glycerol, 0.5% Triton-X, 0.2 mM NaOV_3_, 10 mM NaPO_3_, 10 mM β-glycerophosphate, 10 mM NaF, resuspended in 50 μl 2x SDS-loading buffer (200 mM Tris pH 6.8, 40% glycerol, 4% SDS, 1 M β-mercaptoethanol) and boiled for 10 min at 95°C. For phosphatase treatment, immunoprecipitated beads were incubated with 40 Units of lambda protein phosphate (New England Biolabs) in a 50 μl reaction. Samples were washed once with wash buffer and resuspended in 50 μl 2 x SDS-loading buffer.

### SILAC labeling

To quantify relative phosphorylation on amino acid residues in Mdh1-GFP in *AUP1* and *aup1Δ* backgrounds, cell cultures of each strain were grown separately in SL medium supplemented respectively with 0.003% (w/v) ^13^C_6_ ^15^N_2_ L-lysine (Lys8; Cat. Number 211604102, Silantes, Munich) and 0.001% (w/v) ^13^C_6_^15^N_4_ L-arginine (Arg10; Cat. Number 201603902, Silantes, Munich) or with identical concentrations of unlabeled lysine and arginine, in addition to the standard auxotrophy supplementation. At each time point, 50 OD_600_ units of cells from each labeling regime were combined, washed with cold lysis buffer, and frozen in liquid nitrogen and processed for native immunoprecipitation as described in the previous section. Phosphopeptide ratios were normalized to respective Mdh1-GFP ratios. To follow protein-protein interactions via SILAC-based quantitative MS, we quantified the ratio of proteins co-immunoprecipitated with Mdh1-GFP against the respective signal observed in cells expressing normal, untagged Mdh1. As the signal for untagged cells should be an averaged, constant background, the ratio of this parameter between different genetic backgrounds (*AUP1* vs *aup1Δ*) provides a quantitative assessment of the effects of *aup1* deletion on the different interactions. Briefly cells (PKY392, TVY1 and PKY415, PKY395) were grown as above, with tagged strains (PKY392, PKY415) labeled with heavy isotopes and untagged strains grown in light isotopes (see previous section for details). During cell harvesting, cognate light and heavy labeled cultures were mixed in equal proportions and processed as for native immunoprecipitation, above.

### MS analysis

Mass spectrometric measurements were performed on a Q Exactive Plus mass spectrometer coupled to an EasyLC 1000 (Thermo Fisher Scientific, Bremen, Germany). Prior to analysis phosphopeptides were enriched by TiO_2_, as described previously (64). The MS raw data files were analysed by the MaxQuant software (65) version 1.4.1.2, using a Uniprot *Saccharomyces cerevisiae* database from March 2016 containing common contaminants such as keratins and enzymes used for in-gel digestion.

### Statistical analysis

For detailed statistical methods used in calculating significance of differences in band densitometry and in generating the heat map shown in Figure 1H, please refer to SI appendix.

**For additional methods and experimental details, please refer to SI appendix.**

## Acknowledgements

We wish to thank Chris Meisinger and Corvin Walter for advice on mitochondrial protein phosphorylation, Hillary Voet and Inbar Plaschkes for help with statistical analysis, Einat Zelinger for help with confocal microscopy, Chris Meisinger and Maya Schuldiner for plasmids, and J.C. Martinou for discussions. This work was funded by ISF grant 422/12 (to HA), ISF grant 445/17 (to HA), GIF grant I-111-412.7-2014 (to HA and JD), the Swiss National Science Foundation, grant 310030_184781, and TRANSAUTOPHAGY, COST Action CA15138 (to JD).

## Author contributions

P.K. and V.N. carried out experiments, generated SILAC-labeled samples for MS analysis, and wrote the manuscript. M.R. and J.Z. carried out MS analyses, and analyzed the MS data. J.D. carried out analyses of MS data, planned experiments and wrote the manuscript. H.A. planned experiments and wrote the manuscript.

