## Supplementary information for "Phosphorylation of mitochondrial matrix proteins regulates their selective mitophagic degradation"

**Supplementary information appendix**

**Supplementary Materials and Methods**

**Plasmid construction**

To make plasmids pPKB1 and pPKB2, the *AUP1* reading frame full length and an alternative insert lacking 34 amino acids from the N-terminal of the reading frame, respectively, were amplified using a PCR reaction. The primers P1 (5'-ATACTAACTAGTTACAACATGCGGCTGGGGAATCT ATG-3' ) and P2 (5'-ATACTAATCGATTTAGTTTAATTTTGTTTCGTTAGATTG-3) contain *ClaI* and *SpeI* linkers (SI appendix table III). The PCR product was digested then with *ClaI* and *SpeI* and ligated into a *ClaI* and *SpeI* digested pCU416. To construct plasmid PKB32, the Mdh1-GFP open reading frame (ORF) was amplified by HAY840 yeast genomic DNA using a PCR reaction and oligonucleotides primers P7 Forward: 5'-ATACTAACTAGTTACAACA TGTTGTCAAGAGTAGCTAAAC-3' and P8 Reverse: 5'-ATACTAATCGATGACCTCATACTATACCTG-3' containing *ClaI* and *SpeI* restriction sites (SI appendix table IV). The PCR product was digested with *ClaI*-*SpeI* and ligated into a *ClaI* and *SpeI* digested pCU416, generating a fusion protein under the control of the *CUP1* promoter. Plasmid PKB65 was generated by cloning the Mdh1-GFP reading frame, together with 800 bases of 5' and 500 bases of 3' sequences was amplified using a PCR and primers P5 Forward: 5'-ATACTAGGATCCC AAAAGATCGACGCAATG-3' and P6 Reverse: 5'-ATACTAGAGCTCGACCTCATACTATACCTG-3' (table 4) containing *BamHI* and *SacI* linkers. The PCR product was digested with *BamHI-SacI* and ligated into a *BamHI-SacI* digested pRS415. All the plasmids were verified by sequencing (Hylabs, Rehovot).

**Site directed mutagenesis**

Site directed mutagenesis protocol was carried out according to the Stratagene QuikChange system. *Pfu* DNA Polymerase (Fermentas) was used with a thermocycling protocol followed by removal of the parental strands with *DpnI* digestion. Thermocycling conditions were as follows: Initial denaturation at 95°C for 2 min, then (steps 2-4), 30 sec at 95 °C, 60 sec at 55°C, and 12 min at 68 °C. Steps 2-4 repeated for 18 cycles. PCR products were treated with DpnI (fermentas) for 1 hr at 37°C, and precipitated using with 70% ethanol. DNA pellets were then dried and resuspended with 20 μl TE buffer pH 8. All mutagenesis products were transformed into *E. coli* DH5α, and mutations were verified by sequencing. The oligonucleotide primers used to create site-directed MDH1-GFP variants and combined variants are listed in SI appendix Table III.

**Culture conditions**

Yeast were grown in synthetic dextrose medium (0.67% yeast nitrogen base w/o amino acids (Difco), 2% glucose, auxotrophic requirements and vitamins as required) or in SL medium (0.67% yeast nitrogen base w/o amino acids (Difco), 2% lactate pH 6, 0.1% glucose, auxotrophic requirements and vitamins as required). All culture growth and manipulation were at 26 $^{0}$C. Yeast transformation was according to Gietz and Woods (63).

For nitrogen starvation experiments, cells were grown to mid-log phase ($OD_{600}$ of 0.4-0.6) in synthetic dextrose medium (SD), washed with distilled water, and resuspended in nitrogen starvation medium (0.17 YNB-N (Difco), 2% glucose) for the times indicated in individual experiments. Overexpression studies with *CUP1* promoter-based vectors were carried out by supplementing the medium with 5 μM $CuSO_4$, for both control (empty vector) and overexpressing cells.

**Calculation of statistical significance of differences in % free GFP signals from immunoblots**

Data (% free GFP) were log-transformed before analysis in order to stabilize variances. A repeated measures ANOVA was performed to compare mutants and proteins simultaneously. Thereafter, mutants were compared for each protein and proteins were compared for each mutant by the Tukey HSD test ($p<0.05$). ANOVA analysis was carried out using JMP 12 software.

**Generation of selectivity profile comparisons and heat maps**

To compare between different genotypes, % free GFP values which were recorded from immunoblots of Mdh1-GFP, Aco1-GFP, Aco2-GFP, Qcr2-GFP and Idp1-GFP were normalized, such that each value of % free GFP was divided by the average value of all % free GFP measured for the same protein over all genotypes. This centered the data distribution such that all proteins contributed equally to the correlation calculation. Normalized data points of 3 biological replicates were averaged. We then calculated a Pearson correlation value (r (correlation coefficient) and p – values (significance of correlation) between all phenotype pairs (vector of normalized averaged % free GFP per protein) using the rcorr function in R (Hmisc package). A heat map was generated using the heatmap.2 function in R (gplots package). The heat map cells contains the r correlation measured and colored from negative correlation in red to positive correlation in yellow. The dendogram was generated by calculating the Euclidean distance measure between correlation vectors of each genotype using the complete linkage method.

**Cell fractionation**. Cells (20 $OD_{600}$ units) were collected by centrifugation at 3,500 xg, 5 min, 4°C. The cells were spheroplasted in medium containing 1 M sorbitol and 0.6 mg/ml zymolyase (0.67 % yeast nitrogen base, 2 % glucose, auxotrophic requirements and vitamins as required, 1 M sorbitol, 40 mM HEPES pH 7, 0.6 mg/ml zymolase) for 30 min at 37°C. Spheroplasts were collected by centrifugation at 200xg g for 5 min. The spheroplasts were resuspended on ice in lysis buffer (0.2 M sorbitol, 50 mM potassium acetate, 2 mM EDTA, 40 mM HEPES pH 7, plus protease inhibitors), transferred to a pre-cooled dounce homogenizer and dounced 15 times with a tight fitting pestle. The lysate was then transferred to eppendorf tubes and centrifuged at 300 xg, 4°C, 5 min. 1 ml of cleared supernatant was saved as total extract fraction (T). The rest was transferred to clean eppendorf tubes and centrifuged at 13.000x g, 10 min, 4°C. 1 ml of the supernatant was labeled as S13 (cytosolic proteins). The pellet was labeled as P13. The S13 and total extract fractions (1 ml each) were precipitated with 10% cold TCA by adding 500 µl of 30% TCA, while the P13 fraction was first resuspended in 1 ml lysis buffer and then precipitated with 10 % TCA. For immunoblot analysis, 0.5 $OD_{600}$ equivalents were loaded per lane on SDS-PAGE gels.

**Preparation of whole-cell extracts for western blot analysis**. Cells (10 $OD_{600}$ units) were treated with 10% cold trichloroacetic acid (TCA) and washed three times with cold acetone. The dry cell pellet was then resuspended in 100 µl cracking buffer (50 mM Tris pH 6.8, 6 M urea, 1 mM EDTA, 1% SDS) and vortexed in Tomy MT-360 microtube mixer at maximum speed, with an equal volume of acid-washed glass beads (425-600 µm diameter), for a total of 30 min. Lysates were clarified by centrifugation at 17,000 xg for 5 min and total protein was quantified in the supernatant using the BCA protein assay (Thermo Scientific, Rockford, IL). SDS loading buffer (final concentrations of 100 mM Tris pH 6.8, 20% glycerol, 2% SDS, 500 mM β-mercaptoethanol) was added to the lysate and the samples were warmed to 60°C for 5 min prior to loading on gels. ImageJ software was used for band quantification.

**Immunoblotting**. SDS-10% polyacrylamide gels were transferred to nitrocellulose membrane by using either a wet or semidry blotting transfer system. Membranes were blocked for 1 h with TPBS (137 mM NaCl, 2.7 mM KCL, 10 mM $Na_2HPO_4$, 1.8 mM $KH_2PO_4$, 1 % Tween and 5 % milk powder (BD Difco Skim Milk), followed by incubation with anti-GFP rabbit (1:5000) for 1.5 h, then washed 4x 2 min with TPBS, incubated for 1.5 h with HRP conjugated secondary antibody (1:10000). The membranes were then washed 4x with TPBS, incubated with SuperSignal Chemiluminescence substrate (Thermo Scientific), and exposed to a Molecular Imager ChemiDoc™ XRS imaging system.
